# Cross-Subject EEG-Based Emotion Recognition through Neural Networks with Stratified Normalization

**DOI:** 10.1101/2020.09.18.304501

**Authors:** Javier Fdez, Nicholas Guttenberg, Olaf Witkowski, Antoine Pasquali

## Abstract

Due to a large number of potential applications, a good deal of effort has been recently made towards creating machine learning models that can recognize evoked emotions from one’s physiological recordings. In particular, researchers are investigating the use of EEG as a low-cost, non-invasive method. However, the poor homogeneity of the EEG activity across participants hinders the implementation of any such system by a time-consuming calibration stage. In this study, we introduce a new participant-based feature normalization method, so-called stratified normalization, for training deep neural networks in task of cross-subject emotion classification from EEG signals. The new method is able to subtract inter-participant variability while maintaining the emotion information in the data. We carried out our analysis on the SEED dataset, which contains 62-channel EEG recordings collected from 15 participants while watching film clips. Results demonstrate that networks trained with stratified normalization outperformed standard training with batch normalization significantly. In addition, the highest model performance was achieved when extracting EEG features with the multitaper method, reaching a classification accuracy of 91.6% for two emotion categories (positive and negative) and 79.6% for three (also neutral). This analysis provides us with great insight into the potential benefits that stratified normalization can have when developing any cross-subject model based on EEG.

## Introduction

Emotion recognition has gained great attraction due to its large number of potential applications in fields such as human-computer interaction (e.g., Brave and Nass, 2009), interactive storytelling (e.g., Fels et al., 2011), and mood dis-orders (e.g., El Keshky, 2018). Specifically, researchers are exploiting emotion recognition via EEG signals due to its advantages compared to other low-cost, non-invasive methods such as electromyogram (EMG) and electrocardiography (ECG), whose current limitations restrain them to be used mainly in multimodal emotion recognition (e.g., Dzedzickis et al., 2020), or to facial expression and speech emotion recognition methods, which are susceptible to cognitive bias such as social desirability bias (e.g., Gery et al., 2009; Heuer et al., 2007).

However, the main bottleneck in the development of models trained with EEG signals is the poor homogeneity of between-sessions data and between-participants data, which, interestingly, is not apparent in literature in the context of emotion recognition from facial expressions or other physiological data (Cimtay and Ekmekcioglu, 2020). In order to solve this problem with EEG, current methods rely on participant-dependent models tuned with tedious and time-consuming calibration sessions implemented before each experiment.

In the past years, significant effort has been made in building participant-independent models that eliminate the need for calibration sessions. Specifically, the primary focus of these models is to find common features across participants using algorithms which usually regard variance among individuals as mere statistical noise (Shu et al., 2018). One example is the study of Li et al. (2020), where researchers used an unsupervised deep generative model to capture the emotion-related information between participants. Another example is from Yin et al. (2017), who present an EEG feature selection approach to determine a set of the most robust EEG indicators with stable geometrical distribution across a group of participants. In another study by Li et al. (2018), researchers extracted nine types of time-frequency domain features and nine types of dynamical system features and studied the importance of all those features across different channels, brain regions, rhythms, and features types.

Since the average classification accuracy by selecting robust features is still lower than the participant-dependent models (Shu et al., 2018), researchers are also investigating other approaches such as functional brain connectivity patterns, domain adaptation, or hybrid methods. An example of cross-subject functional brain connectivity investigation is from Cao et al. (2020), who studied the key information flow of the different parts of the brain with minimum spanning trees (MST). About domain adaption, the study by Chai et al. (2016) presents several unsupervised domain adaptation techniques based on autoencoders for non-stationary EEG-based emotion recognition. Furthermore, Cimtay and Ekmekcioglu (2020) analyzes the use of pre-trained convolutional neural network (CNN) architectures to improve the feature extraction and inherent exploitation of the domain adaptation. Lastly, the report by Yang et al. (2019) gives an example of a hybrid method for cross-subject emotion recognition by extracting multiple features for the formation of high-dimensional feature space.

Another approach, which is more relevant to this work, is the normalization of the data to improve the cross-subject emotion recognition. The studies by Koelstra et al. (2012) and Jatupaiboon et al. (2013) already gave the first insights into the advantages of this approach after applying participant-based data normalization to reduce the inter-participant variability. Later on, the work of Arevalillo-Herráez et al. (2019) exploited this result and proposed a nonlinear data transformation that seamlessly integrated individual traits into an inter-participant approach.

Throughout this article, we extend those observed results by introducing and assessing a participant-based feature normalization approach, so-called *stratified normalization*. In comparison and to deepen the evaluation, we also present the results for neural networks trained with standard batch normalization under different labeling and feature extraction methods.

To encourage further research on these topics, we have made the source code of this work freely accessible to all ^1^.

## Materials and methods

We designed an experiment to capture the effects of one control variable – the experimental data – and three independent variables – the labeling method, the feature extraction method, and the normalization method – onto the dependent variable that we aim to assess – the cross-subject emotion recognition accuracy of the trained models. Figure 1 details the variables and their conditions for the experiment. The four sections of this chapter detail each of the experiment’s control and independent variables, respectively.

**Fig. 1.**
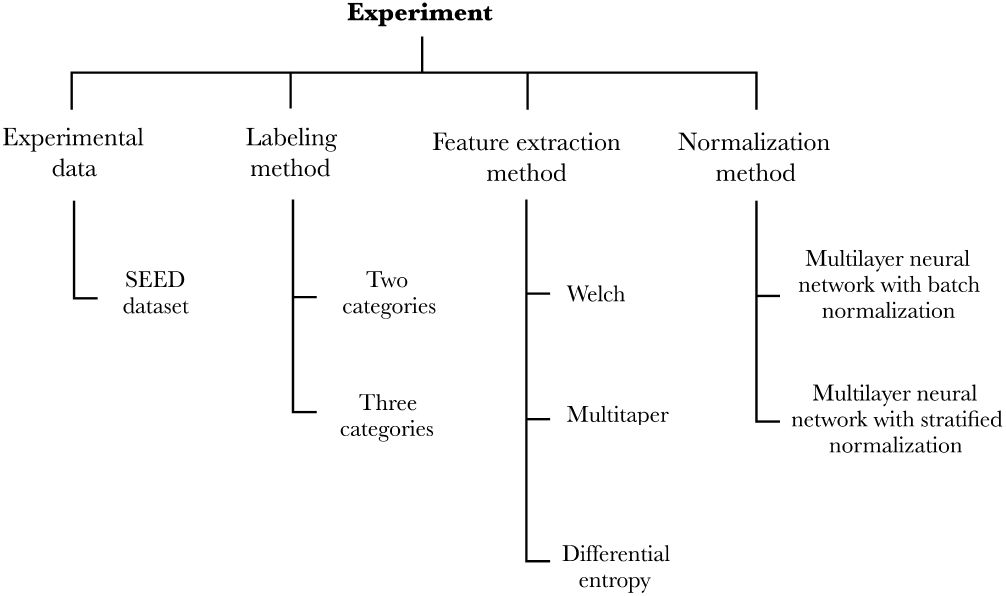
Our experimental setup captures the effects that the experimental data, labeling method, feature extraction method, and normalization method have onto the emotion recognition accuracy of the models.

Our methodology unfolds as follows. Firstly, we picked an EEG dataset and a preprocessing stage of the data so as to feed our models (c.f., A. Experimental data). Secondly, we prepared two strains of models to perform either binary or ternary classification (c.f., B. Labeling method). Subsequently, we defined models that would extract the features according to the three conditions of the feature extraction method variable (c.f., C. Feature extraction method). Lastly, we implemented the two conditions of the normalization method variable (c.f., D. Normalization method) and assessed each model (2 × 3 × 2 = 12 models in total) through a leave-one-out cross-validation over all participants from the EEG dataset.

### A. Experimental data

Despite the large number of potential applications of emotion recognition from EEG signals, to the best of our knowledge, MAHNOB-HCI (Soleymani et al., 2012), SEED (Duan et al., 2013a; Zheng and Lu, 2015) SEED IV (Zheng et al., 2019), and DEAP (Koelstra et al., 2012) are the only four publicly available emotional EEG datasets on the topic. From those four datasets, we chose to work with the SEED dataset because (1) the collected data is well balanced between sessions and participants, and (2) the literature offers many reports on cross-subject emotion recognition models using this dataset.

The SEED dataset contains 62-channel EEG data collected from 15 participants, who carried out three sessions over the same 15 film clips. An emotional rating was previously assigned to each film clip and obtained by averaging the ratings of 20 participants who were asked to indicate one keyword (positive, neutral, or negative) after watching them. In the subsequent experiment with EEG recordings, the clips used across sessions and participants at each trial shared the same pseudo-random order of ratings to smoothly balance evoked emotions throughout the EEG recordings.

In addition to the raw EEG data, the SEED dataset contains a preprocessed version of the signals, consisting of a downsampling to 200 Hz and a noise filtering with a bandpass filter of 0.5-70 Hz. Since the downsampling reduces the high dimensionality and the noise filtering increases the signal-to-noise ratio (Bigdely-Shamlo et al., 2015), we have chosen to use this preprocessed data for our analysis.

### B. Labeling method

Our first independent variable indicated the number of emotional classes: two categories (positive and negative) and three categories (positive, neutral, and negative). The two approaches can be conveniently compared due to the large number of articles that reported their results with binary classification (e.g., Li et al., 2018; Yang et al., 2019; Li et al., 2020; Cimtay and Ekmekcioglu, 2020) or ternary classification (e.g., Chai et al., 2017; Zhang et al., 2019; Lan et al., 2019; Cimtay and Ekmekcioglu, 2020).

### C. Feature extraction method

The second independent variable concerns the feature extraction method, which refers to either the (1) Welch, (2) multitaper, or (3) Differential Entropy (DE) method in our study. The first two methods were selected since they belong to the Power Spectral Density (PSD) category, which, according to Craik et al. (2019), is a typical approach when training deep neural networks for EEG classification tasks. Our selection of DE was based on the high accuracy reported in some EEG emotion recognition studies, such as by Duan et al. (2013a) and by Chen et al. (2019).

For all three methods, the total number of extracted features for each trial is 248 (62 channels x 4 band frequencies). The four band powers for each EEG signal correspond to the theta rhythm (4-7 Hz), alpha rhythm (8-13 Hz), beta rhythm (14-30 Hz), and gamma rhythm (31-50 Hz). The delta rhythm (0.5-4 Hz) was excluded as it is traditionally associated with sleep stages (de Andrés et al., 2011) and therefore assumed to be less relevant to our study.

#### C.1. Welch’s method

Welch’s method (Welch, 1975) is an approach to estimate the power of a signal at different frequencies. It is carried out by averaging consecutive periodograms of small time-windows over the signal. To encompass at least two full cycles of the lowest frequency of interest (4 Hz), the duration of the time-windows was set at 0.5 seconds, with an overlap of 0.25 seconds between each consecutive window. To smooth the discretization process, each window was filtered with a Hann function. The band frequencies were thereafter extracted from the PSD by implementing Simpson’s rule, which approximates integrals using quadratics.

#### C.2. Multitaper method

Multitaper method is an alternative to Welch’s method, which still produces high variance for the direct spectral estimation (Mansouri and Castillo-Guerra, 2019). This method reduces the variance of the spectral estimation by using multiple time-domain windows rather than a single-domain window. As well as Welch’s method, the band frequencies were extracted by implementing Simpson’s rule over the PSD.

#### C.3. Differential entropy method

The DE is used to measure the complexity of a continuous random variable (Duan et al., 2013b). Its calculation formula can be expressed as,

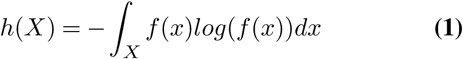

where X is a random variable and f(x) is the probability density function of X. When the time series X obeys the Gaussian distribution N(*µ, σ*^2^), its differential entropy can be defined as,

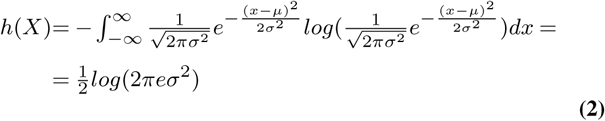

For a fixed-length EEG sequence, we can approximate the EEG data to a Gaussian distribution, constructing features using equation 2.

### D. Normalization method

The last independent variable is the normalization method, which indicates the two types of classifiers implemented in this study: multilayer neural network with batch normalization and multilayer neural network with stratified normalization.

The models used to classify emotions are independent of the normalization method, meaning that batch and stratified normalizations apply to the same neural architectures (c.f., Figure 2). Both types of classifiers were trained for 100 epochs; the whole training data (15 participants x 3 sessions x 15 trials) was input in one unique batch; the learning rate was set to 0.005 for the first 40 epochs, then decreased to 0.001 for the remaining 60 epochs; the optimizer was the Adam optimizer (Kingma and Ba, 2014); the loss function was the negative log-likelihood loss (Zhu et al., 2018). For further information about the hyperparameters or architecture of the neural network, please refer to the source code.

**Fig. 2.**
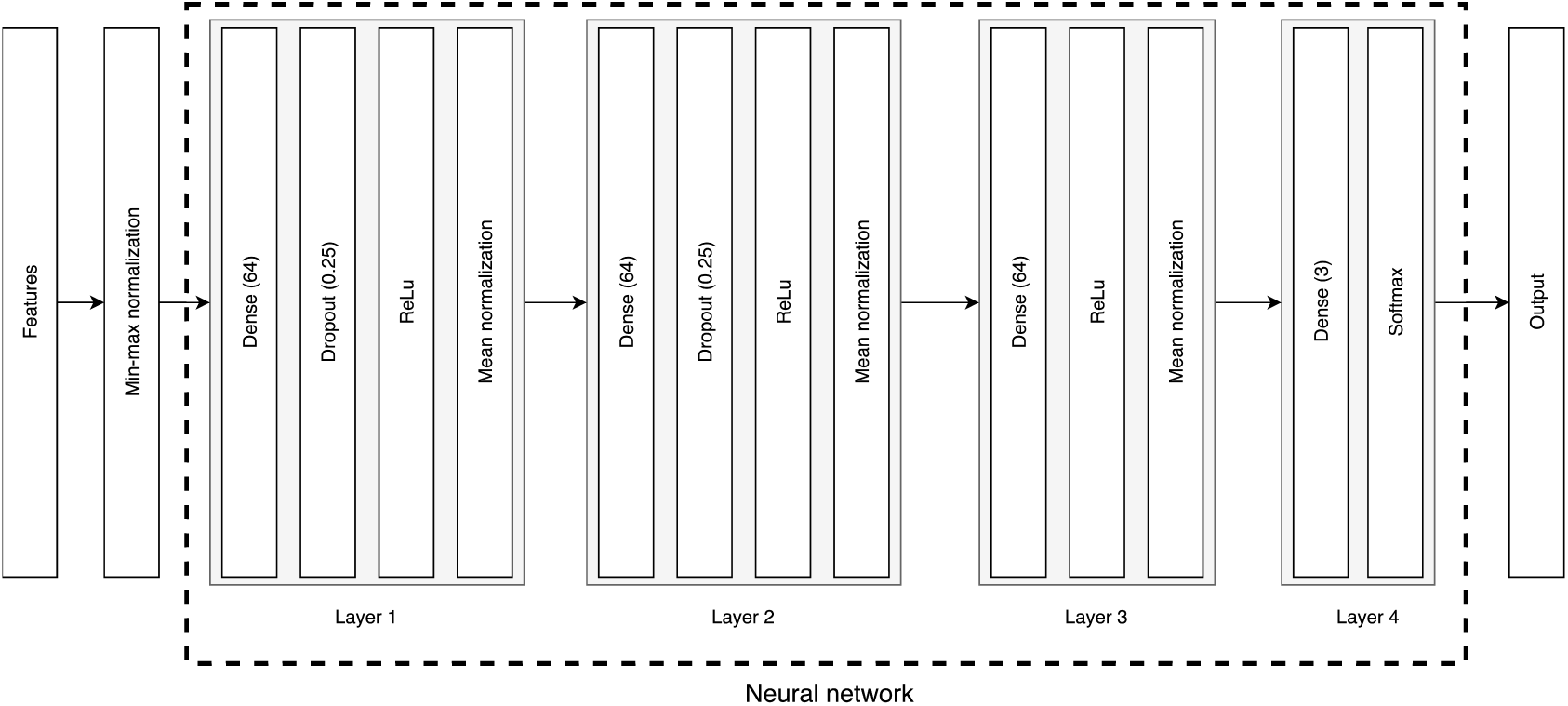
Architecture of the classifiers. Features first go through a min-max normalization of the data before being input to the neural network. The first three layers consist of dense layers with 64 neurons, a dropout for the first two, ReLu activations, and either a batch or a stratified normalization. The last layer is a 3-neuron dense layer that outputs the classification prediction through a Softmax function.

#### D.1. Batch normalization

Batch normalization was first introduced by Ioffe and Szegedy (2015) in order to address the problem of *internal covariate shift*, an unwanted drift in the distribution of neuron’s activations resulting from the learning process. As explained further below, we slightly adapted the method for our purposes. Figure 3 illustrates our implementation of the batch normalization method.

**Fig. 3.**
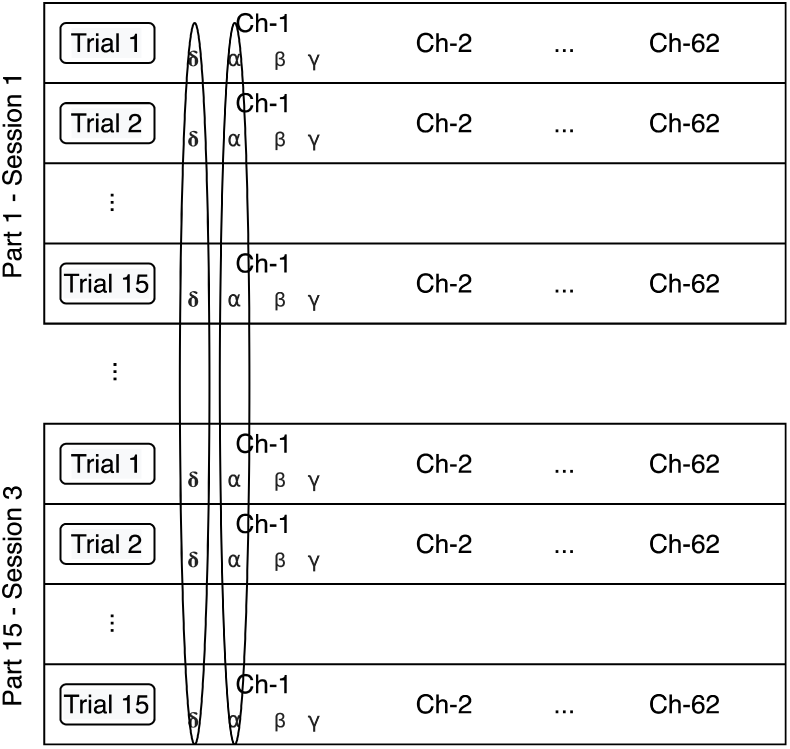
Batch normalization method. The data is normalized per feature, independently of the participant and session.

The extracted features are first min-max normalized according to the following equation:

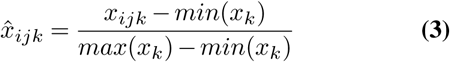

where the parameter *i* indicates the number of the participant and session (45 in total), *j* refers to the number of the trial (15 in total), and *k* identifies the number of the feature (248 in total).

The output of this min-max normalization is input to the neural network, which implements mean normalization of the output of each of the first three layers according to equations 4, 5 and 6.

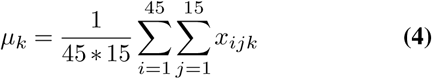

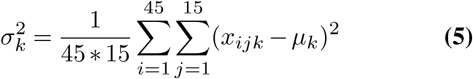

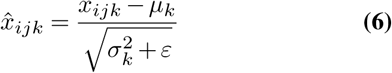

These equations correspond to the first three steps of the batch normalization transform described in Ioffe and Szegedy (2015). The fourth step of the algorithm is a scale and shift of the normalized values, where the parameters are learned along with the original model parameters. However, we observed that this step decreased the emotion recognition accuracy, so we decided to exclude it from our analysis.

#### D.2. Stratified normalization

The stratified normalization consists of a feature normalization per participant and session. Figure 4 details our implementation of the stratified normalization method.

**Fig. 4.**
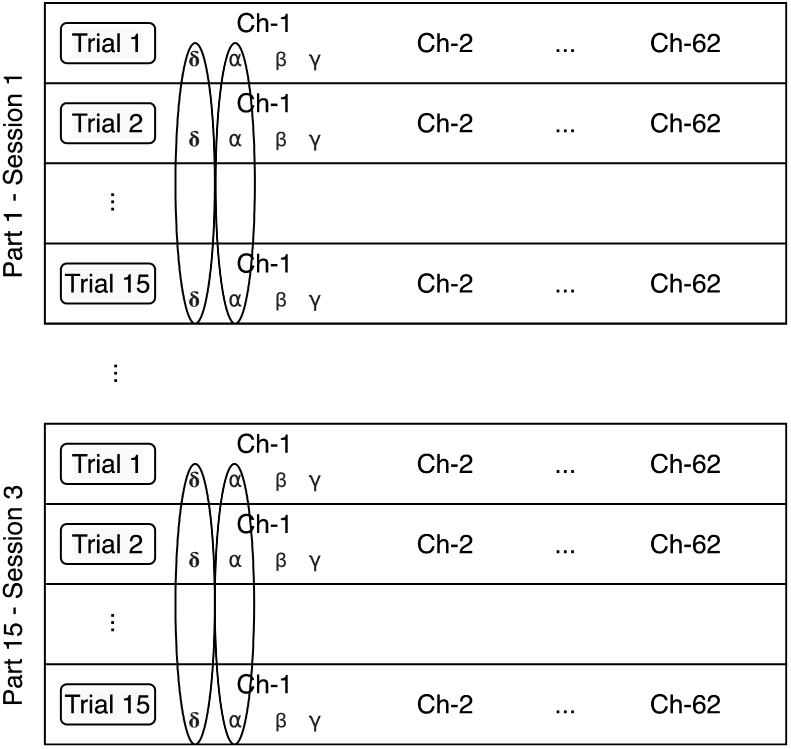
Stratified normalization method. The data is normalized per feature, participant and session.

The extracted features are first min-max normalized according to the following equation:

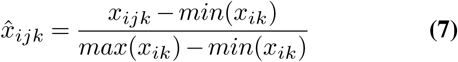

The output of this min-max normalization is input to the neural network, which implements mean normalization at the output of each of the first three layers according to equations 8, 9 and 10.

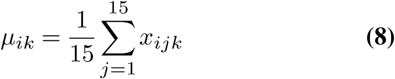

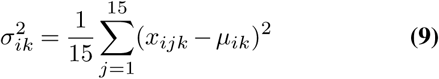

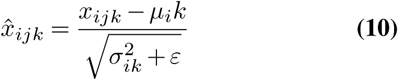

## Results and discussion

This section first presents the results of the experiment, then analyzes the between-participant variance and cross-subject emotion recognition in the layers of the neural networks, and finally compares the results of this work with state-of-art literature.

### A. Overall evaluation

Figure 5 reports the performance, tested after 100 epochs of training, of the models in each experimental condition.

**Fig. 5.**
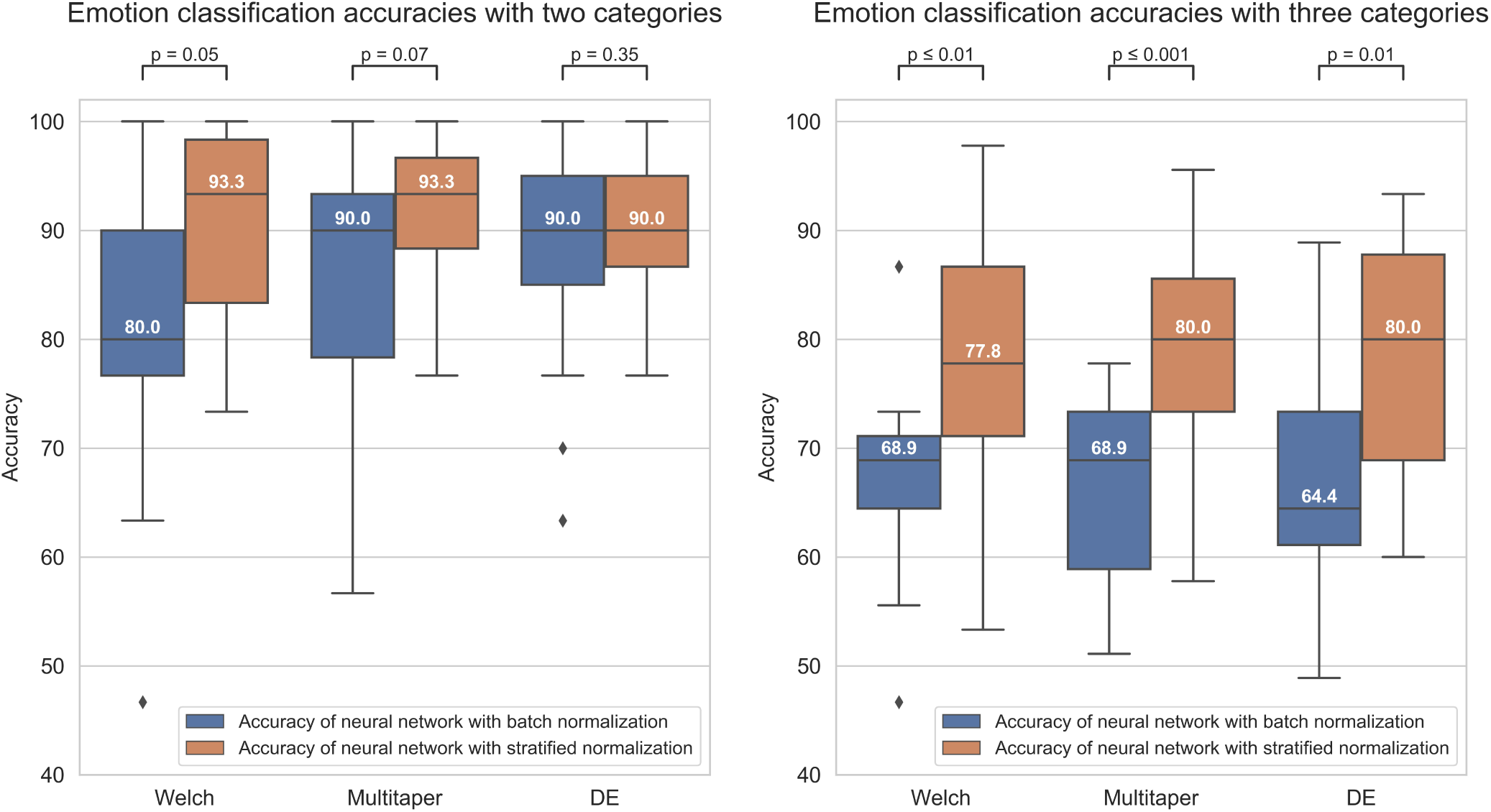
The boxplots of the figure indicate the distribution of the emotion recognition accuracies (y-axis) of the models in each experimental condition, with either (1) a binary classification (left graph) or ternary classification (right graph) labeling method, (2) a Welch, multitaper, or Differential Entropy (DE) feature extraction method (graph’s columns), (3) a batch (blue bars) or stratified (orange bars) normalization method. The value inside the boxplots is the median value of the distribution. Besides, the figure also reports the p-value of the independent-samples t-test between the two normalization method conditions for each feature extraction method condition.

To evaluate the performance of the models, we ran a three-way ANOVA where the between factors were the (1) labeling, (2) normalization, and (3) feature extraction methods. The statistical results revealed an effect of the labeling method (F(1, 168) = 97.7, p < 0.001, 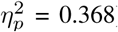), meeting our expectations of an overall better performance in task of binary classification compared to ternary classification. A strong effect of the normalization method was captured as well (F(1, 168) = 33.8, p < 0.001, 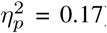), confirming that stratified normalization is a more efficient approach than batch normalization in such an experimental context. However, our manipulation of the feature extraction method did not elicit any effect (F(2, 168) = 0.32, p = 0.73, 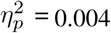), meaning that all methods have the potential to perform equally well. No two-way or three-way interactions were captured by our analysis either (Fs < 4).

Despite a lack of effect of the feature extraction method, we’ve decided to look deeper into the performance of models according to each feature extraction method. Our aim was to allow comparison with the state-of-the-art literature and to further deepen our theoretical interpretation of the results. For binary classification, the methods that performed with the highest accuracy for batch and stratified normalization were DE (M = 0.876, SD = 0.101) and multitaper (M = 0.916, SD = 0.074), respectively. For ternary classification, Welch (M = 0.671, SD = 0.088) and multitaper (M = 0.796, SD = 0.104) were the optimal methods for batch and stratified normalization, respectively. Thus, models using the multitaper feature extraction method in combination with stratified normalization elicit the highest performance both in binary and ternary classification tasks. We will therefore focus on these models for the remaining of our study.

### B. Descriptive summary for the multitaper and stratified methods

In this section, we present a descriptive summary of the results obtained when using, as normalization method, the stratified normalization and, as feature extraction method, the multitaper method. We selected the former method since it resulted in being statistically significant compared to batch normalization in terms of models’ accuracy. About the feature extraction method, the statistical test did not draw any significant result that could allow us to conclude on which method is the most effective. However, we decided to select the multitaper method since it was the feature extraction method with which models performed with the highest accuracy on average, for both binary (M=0.916, SD=0.074) and ternary classification (M = 0.796, SD = 0.104).

Table 1 indicates the leave-one-out classification accuracies for models based on the multitaper and stratified normalization methods. Each column represents the test accuracy of the models on the untrained data (1 participant out of 15) throughout our 15-fold cross-validation design. The comparison between binary and ternary classification results highlights a moderate correlation between the models’ performance on these two tasks (Pearson correlation of 0.343), suggesting that models extract participant identification information with some consistency over tasks (but see C. Evaluation on input, hidden and output layers of the neural network).

**Table 1.** Leave-one-out classification accuracies for two and three categories. Participant numbers (s01-s15) correspond to each test data of the 15-fold cross-validation.

| Participant No. | s01 | s02 | s03 | s04 | s05 | s06 | s07 | s08 | s09 | s10 | s11 | s12 | s13 | s14 | s15 |
| --- | --- | --- | --- | --- | --- | --- | --- | --- | --- | --- | --- | --- | --- | --- | --- |
| Pos-Neg | 76.7 | 80.0 | 93.3 | 86.7 | 96.7 | 90.0 | 93.3 | 100.0 | 96.7 | 96.7 | 100.0 | 80.0 | 93.3 | 90.0 | 100.0 |
| Pos-Neu-Neg | 77.8 | 57.8 | 93.3 | 82.2 | 84.4 | 80.0 | 80.0 | 91.1 | 84.4 | 71.1 | 75.6 | 86.7 | 62.2 | 71.1 | 95.6 |

The confusion matrices for the multitaper method when combining the results of the leave-one-out cross-validation of all the participants for two and three categories are shown in Tables 2 and 3, respectively. The classification accuracy for positive (92.44%) and negative (90.67%) labels is well balanced in binary classification. However, for ternary classification, while the classifiers keep an accurate performance for positive labels (90.67%), their accuracy is lowered for negative (75.11%) and neutral (78.34%) labels, indicating that introducing the neutral labels hinders the classification of the negative labels.

**Table 2.**
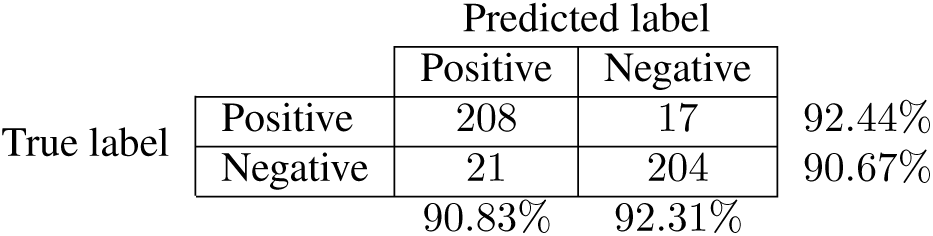
Confusion matrix when combining the results of the leave-one-out cross-validation of all the participants for two categories.

**Table 3.**
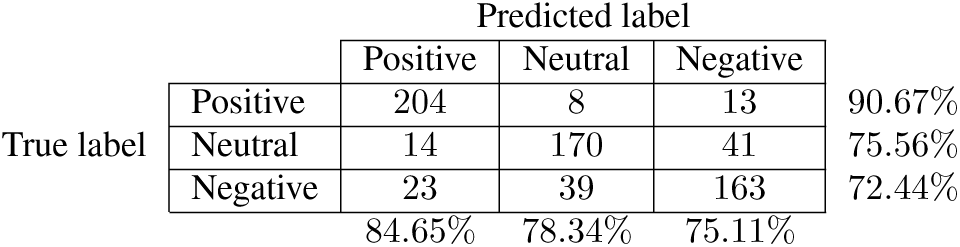
Confusion matrix when combining the results of the leave-one-out cross-validation of all the participants for three categories.

Table 4 lists a performance comparison between state-of-the-art models and our proposed method for two categories (positive and negative) and three categories (positive, neutral, and negative).

**Table 4.** Performance comparison between this work and the state-of-art literature.

| Reference | SEED performance |  |
| --- | --- | --- |
|  | Acc (Pos-Neg) | Acc (Pos-Neu-Neg) |
| Chai et al. (2017) | - | 80.4 |
| Li et al. (2018) | 83.33 | - |
| Zhang et al. (2019) | - | 82.1 |
| Lan et al. (2019) | - | 72.47 |
| Yang et al. (2019) | 89.0 | - |
| Li et al. (2020) | 85.81 | - |
| Cimtay and Ekmekcioglu (2020) | 86.5 | 78.3 |
| This work | 91.6 | 79.6 |

For binary classification, the last benchmark was reported by Yang et al. (2019), who themselves obtained an accuracy of 89.0% by first extracting multiple features for the formation of high-dimensional features and then integrating the significance test/sequential backward selection with the support vector machine for the classification. Our best accuracy for binary classification is 91.6%, which overpasses all reported methods.

For ternary classification, our best accuracy is 79.6%. From the articles listed, our proposed method is overpassed by Zhang et al. (2019), whose model based on convolutional neural network (CNN) and deep domain confusion (DDC) achieved an accuracy of 82.1%, and by Chai et al. (2017), who reached 80.4% by using adaptive subspace feature matching (ASFM). Nevertheless, compared with our approach, both Zhang et al. (2019) and Chai et al. (2017) used the validation set during the training process, which can increase the cross-subject accuracy.

### C. Evaluation on input, hidden and output layers of the neural network

In the previous sections, we evaluated the normalization methods and established that stratified normalization improves the cross-subject emotion recognition accuracy. Nevertheless, an interesting question to be asked is at what depth of the neural network does stratified normalization help increase the accuracy. To answer this question, we further analyzed the emotion recognition accuracy of the models, and their ability to capture and exploit the *brain signature* of each participant – defined as the part of information extracted from the brain signals that is specific to that participant (also called *subject-related component* by Arevalillo-Herráez et al. (2019)), such that *it can directly inform us on which participant it’s been extracted from*. Intuitively, we would expect the emotion recognition accuracy to increase with each layer of the neural network, while the brain signature would fade out due to a decrease of the inter-participant variability with each data normalization.

Therefore, we carried out the evaluation in accordance with the following methodology. For starters, we retrained our models in a 3-fold cross-validation design, using the data of 10 participants for training and 5 for testing. For each of the 3 testing sessions, we recorded the output, or *predicted values*, of each of the model’s layers (after normalization for the input, first hidden, second hidden, and third hidden layers, and after softmax for the output layers). We then fed these predicted values to a series of Support Vector Machines (SVM) with RBF kernels (Chang et al., 2010). To do so, we mixed the data of all 5 test participants and ran a 5-fold cross-validation per layer on the following tasks. In one case, we used the emotional ratings of the 5 test participants as labels, thus evaluating the capacity of each layer to contribute to the emotion recognition accuracy of the model. In another case, we used as labels the participant identification numbers (from 1 to 5), here evaluating the amount of brain signature still available at each layer. The classification results are shown in Figure 6. We also separated the dataset between two and three emotional categories for running the analyses.

**Fig. 6.**
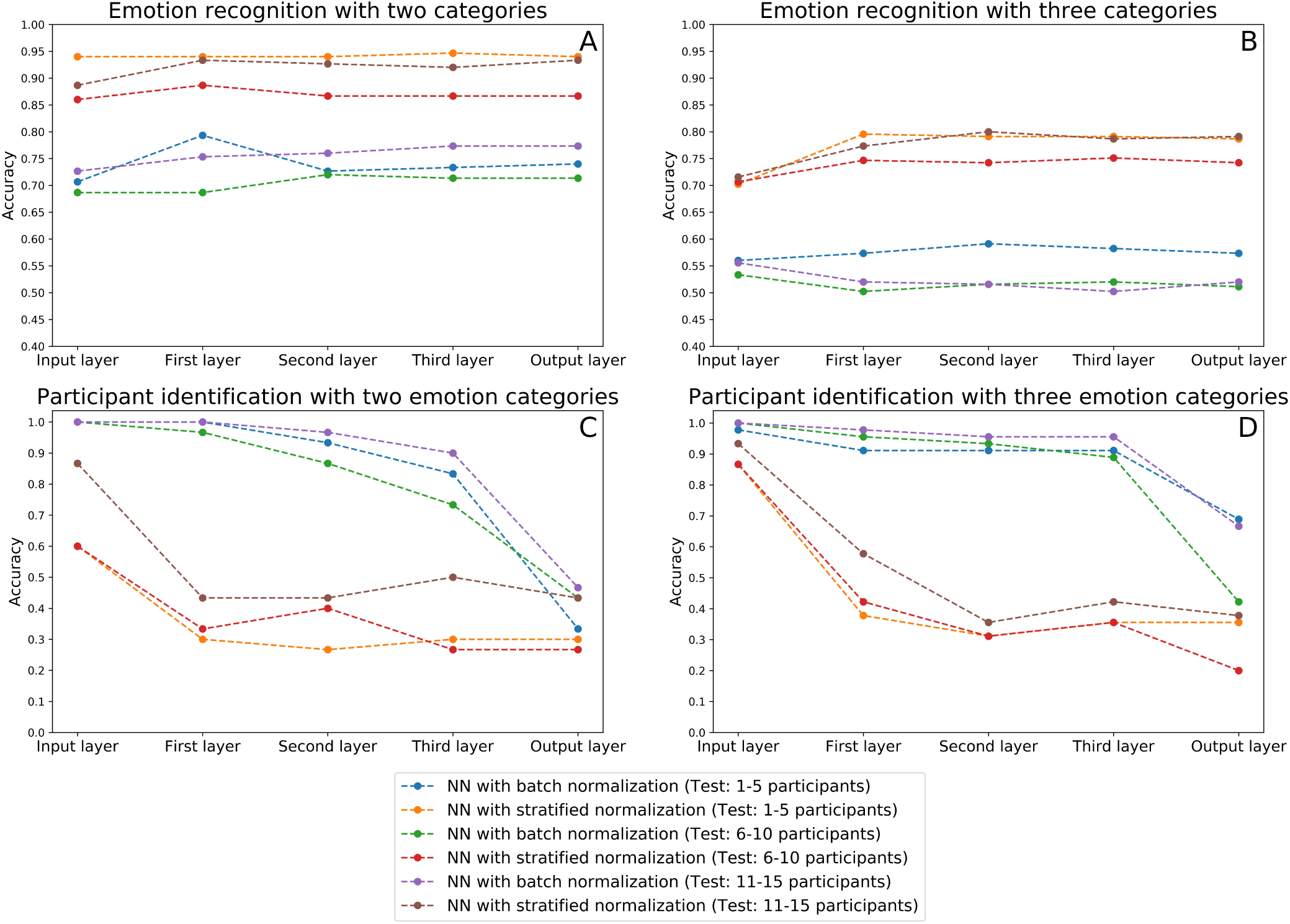
This figure indicates the classification results for the emotion recognition and participant identification in the input, hidden layers, and output of the neural network using multitaper as feature extraction method. Figure 6A and Figure 6B plot the emotion recognition accuracies, and Figure 6C and Figure 6D show the participant identification accuracy.

The statistical analysis of the results is carried out in the following subsections. The dependent variable to study is the classification accuracy obtained. The between factors are the (1) labeling method (two categories, three categories), (2) normalization method (batch or stratified normalization), and (3) layer’s depth (input layer, first layer, second layer, third layer, output layer).

#### C.1. Emotion recognition

To evaluate the emotion recognition in the layers of the neural network, we first implemented a three-way ANOVA test. Results indicated no effect of the layer’s depth (F(4, 40) = 1.49, p = 0.22, 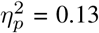). However, we were able to capture main effects of labeling (F(1, 40) = 401.4, p < 0.001, 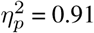) and of normalization (F(1, 40) = 541.0, p < 0.001, 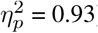) methods. Besides, results indicated that there was an interaction between the labeling method and the normalization method (F(1, 40) = 7.38, p = 0.010, 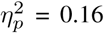). Hence, we implemented a two-way ANOVA test for each of the two conditions of the labeling method. For both of them, the normalization method was found statistically significant (F(1, 20) = 180.5, p < 0.001, 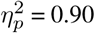 for two categories and F(1, 20) = 405.9, p < 0.001, 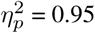 for three categories).

Although the ANOVA didn’t capture any other interaction with layer’s depth (all Fs < 2), we also evaluated normalization methods separately at the input and output layers of the neural network. Results of two-way ANOVA tests for the input layer (F(1, 8) = 156.8, p < 0.001, 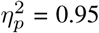) and the output layer (F(1, 8) = 114.1, p < 0.001, 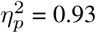) both indicated that stratified normalization surpasses batch normalization.

Therefore, (1) the emotion recognition accuracy does not increase significantly along with the layers of the neural network, and (2) the stratified normalization outperforms batch normalization in emotion recognition, as we already concluded above.

#### C.2. Participant identification

To analyze results in terms of the participant identification accuracy, we first ran a three-way ANOVA test, which pointed out an interaction between the three factors (F(4, 40) = 3.44, p = 0.017, 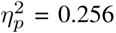), and main effects of the labeling method (F(1, 40) = 6.15, p < 0.05, 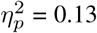), of the normalization method (F(1, 40) = 401.1, p < 0.001, 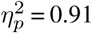) and of the layer’s depth (F(4, 40) = 57.27, p < 0.001, 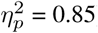). As a result, we carried out a two-way ANOVA test for each condition of the normalization method separately.

For the batch normalization method, we found a significant interaction between the labeling method and the layer’s depth (F(4, 20) = 3.07, p = 0.040, 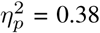). Hence, we implemented a one-way ANOVA test for each condition of the labeling method, each revealing effects of the layer’s depth (F(4, 10) = 60.3s, p < 0.001, 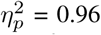 for two categories and F(4, 10) = 15.7, p < 0.001, 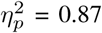 for three categories). Specifically, we found out by implementing several independent-samples t-tests between each pair of consecutive layers that the accuracy only drops significantly between the third and the output layers (t(4) = 6.54, p = 0.003, d = 7.70 for two categories and t(4) = 3.72, p = 0.043, d = 10.7 for three categories).

For the stratified normalization method, analyses showed a significant effect of the layer’s depth (F(4, 20) = 26.9, p < 0.001, 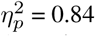). Subsequently, we implemented a two-way ANOVA test between each pair of consecutive layers and this time, we captured a significant drop in accuracy between the input and the first layer (F(1, 8) = 43.6, p < 0.001, 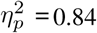). This result suggests a strong correlation between the use of the stratified normalization method and an early drop in participant identification accuracy.

Lastly, to determine if the normalization methods were statistically different, we computed a two-way ANOVA test for each layer of the neural network. All analyses revealed statistically significant effects of the normalization method (F(1, 8) = 20.4, p = 0.002, 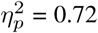 for the input layer, F(1, 8) = 217.6, p < 0.001, 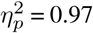 for the first layer, F(1, 8) = 352.1, p < 0.001, 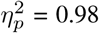 for the second layer, F(1, 8) = 118.9, p < 0.001, 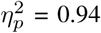 for the third layer, F(1, 8) = 8.83, p = 0.018, 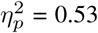 for the output layer). This confirms that the normalization method shows an effect for every layer’s depths, including for the input and the output layers.

Hence, (1) both normalization methods significantly reduce the participant identification information, (2) the layers where the participant identification information is significantly reduced varies depending on the normalization method (output layer for batch normalization and first layer for stratified normalization), and (3) the stratified normalization over-passes batch normalization, having an accuracy of M = 0.33, SD = 0.072 for two categories and M = 0.31, SD = 0.079 for three categories in the last layer, where the chance level is 0.2 since we are classifying five participants.

To conclude, as hypothesized, the decrease of participant identification accuracy observed for both normalization methods confirms that the brain signature is effectively suppressed throughout the Neural Network, preventing the SVM classifiers from recognizing which participant their input data belongs to. Still, we can still see that some participant identification information remains in the output of the models. Indeed, if there wasn’t, then the accuracy of the participant identification would be at a chance level of 20% (considering 5 participants), but instead, it is still at 33% (M = 0.33, SD = 0.072) for two categories and 31% (M = 0.31, SD = 0.079) for three categories in the last layer of the models with stratified normalization. This difference is much higher with batch normalization, 41% (M=0.41, SD=0.057) for two categories and 59% (M=0.59, SD=0.012) for three categories.

We are now wondering about the type of mechanism that operates the brain signature suppression in the model with stratified normalization, and specifically about how it affects the data.

#### C.3. Visualization on the input and output layers of the neural network

To have a clearer visualization of the classifier’s performance, we run the dimensionality reduction tool UMAP (McInnes et al., 2018) to reduce to two dimensions all the predicted values for the input and output layers of the neural network.

Figure 7 and Figure 8 show the embedding of the predicted values at the input layer of the neural network. We have established in the section above that a significant amount of brain signature is already suppressed by this layer. Interestingly, the UMAP for batch normalization shows a handful of compact clusters. These clusters match relatively well the participant numbers, but not the emotional rating. Conversely, the UMAP for the stratified normalization is much more spread out along both dimensions, and neither the emotion ratings nor the participant numbers are recognizable in the cloud of embeddings. This difference suggests that the stratified normalization method induces a redistribution of the activations in output of the first layer in a more spread space of representation, possibly to facilitate further processing in the rest of the network. Then the spread would cause the embeddings to lose their information about the participant number.

**Fig. 7.**
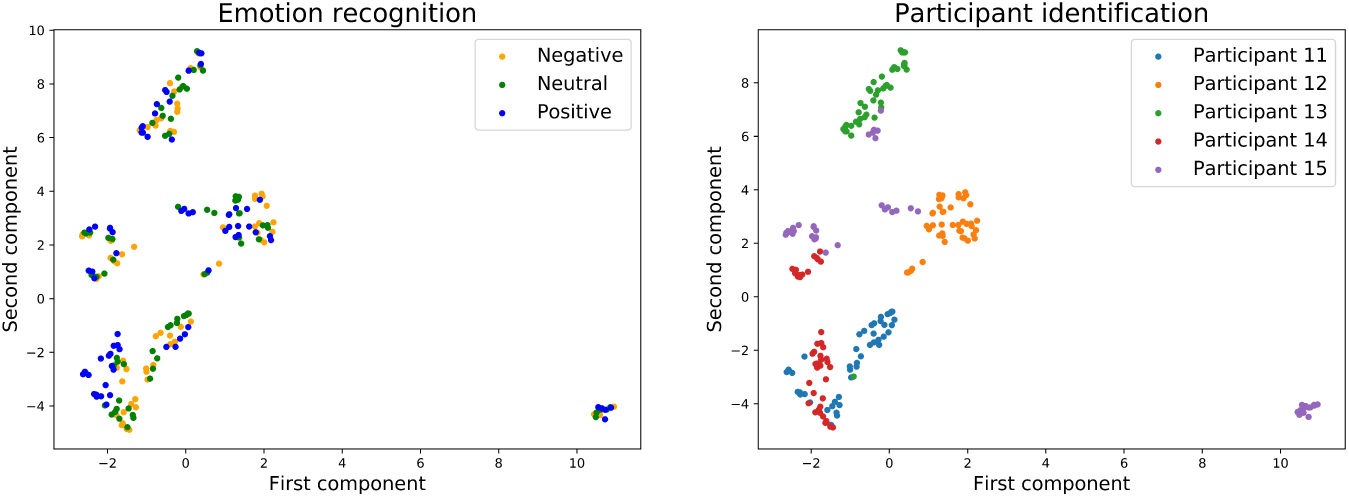
Embedding of the input of the neural network with batch normalization and with three emotion categories.

**Fig. 8.**
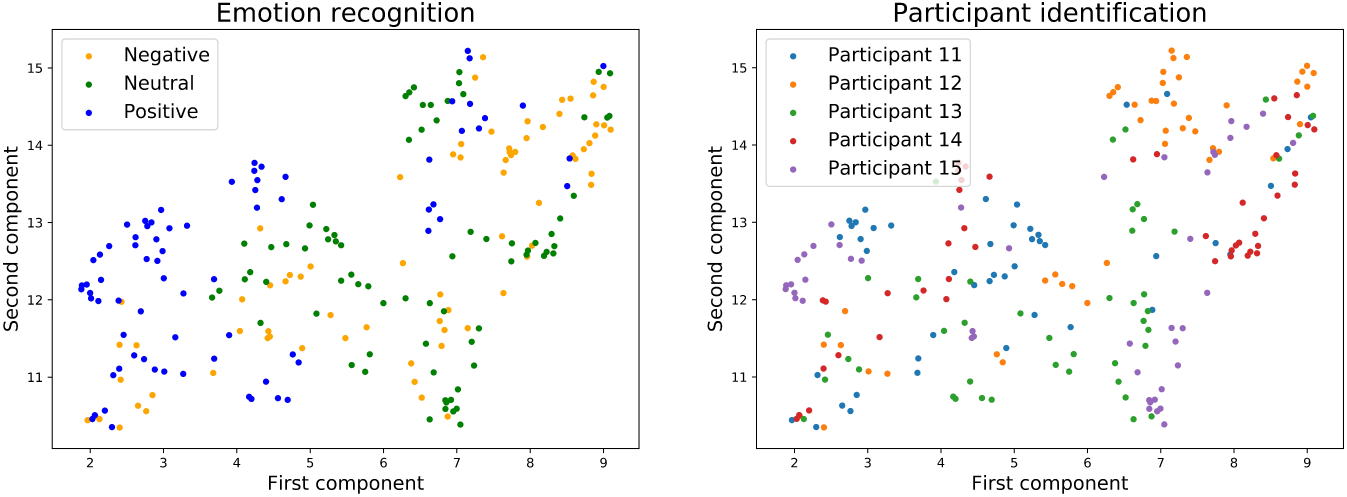
Embedding of the input of the neural network with stratified normalization and with three emotion categories.

Figure 9 and Figure 10 show the embedding of the predicted values at the output layer of the neural network. Our previous results showed that at the output layer, the emotion recognition accuracy is higher and the participant identification accuracy is lower for models trained with stratified normalization. Indeed, this time, embeddings are more compact on the UMAP for the stratified normalization rather than for the batch normalization, and easily recognizable for the emotion ratings rather than for the participant numbers – for which the spread of colors seems to indicate that most of the brain signature is gone indeed.

**Fig. 9.**
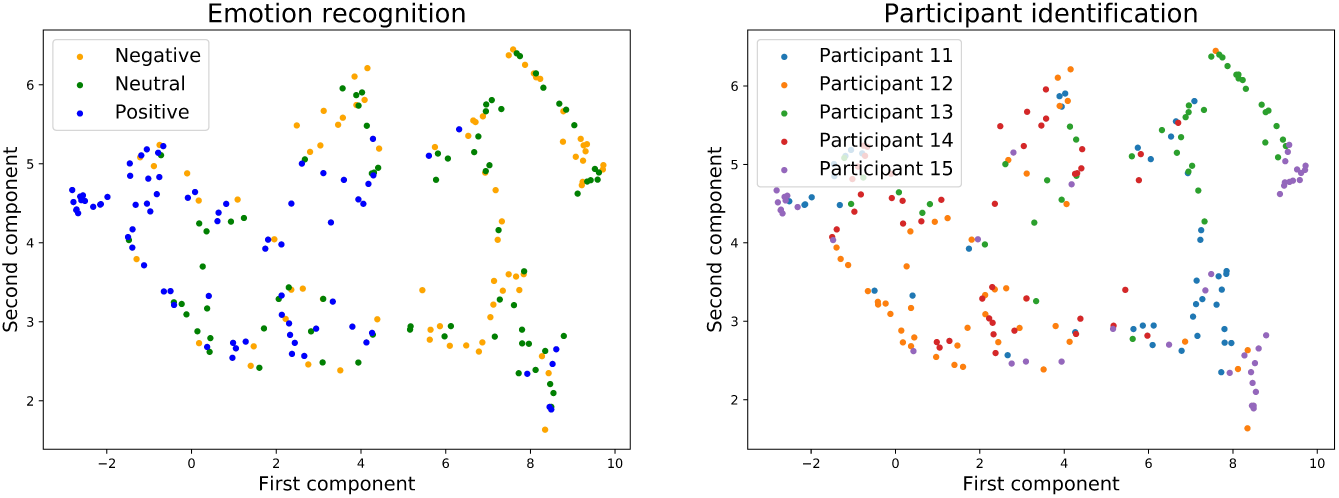
Embedding of the output of the neural network with batch normalization and with three emotion categories.

**Fig. 10.**
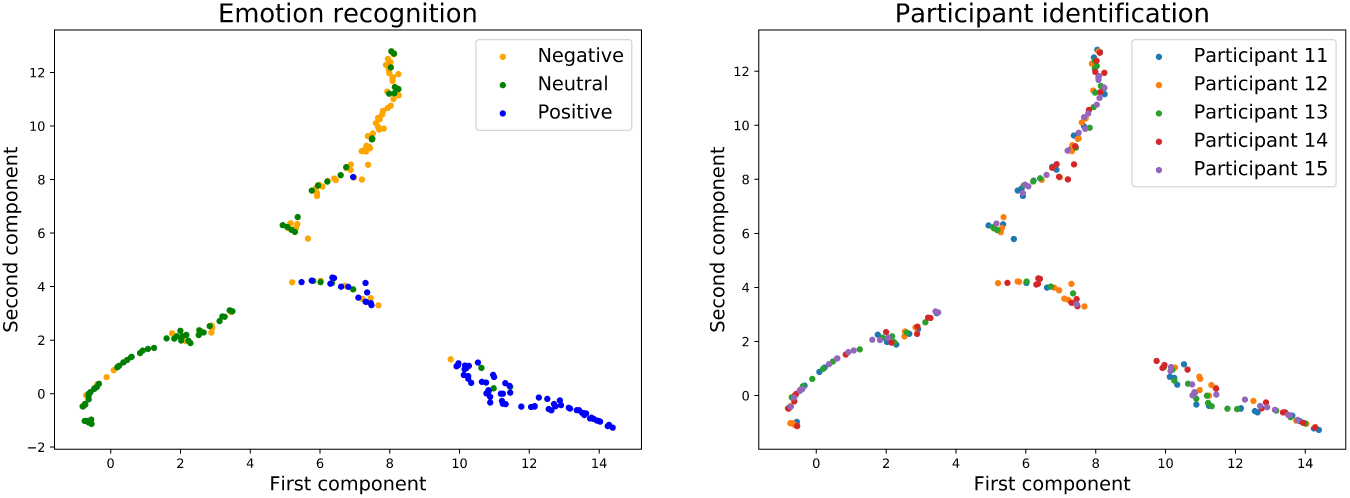
Embedding of the output of the neural network with stratified normalization and with three emotion categories.

## Conclusion

In recent years, researchers have introduced and evaluated different approaches that permit to build robust participant-independent models without the need for prior recorded data from each participant. Specifically, the primary focus is on finding features that do not vary across participants. However, since these methods still present lower accuracy than participant-dependent models, researchers are also investigating other approaches, such as data normalization. In this study, we propose and evaluate a new participant-based feature normalization method, so-called *stratified normalization*, to improve the cross-subject emotion recognition accuracy of participant-independent models.

The evaluation of this method has been carried out by setting an experiment where we recorded the effects of three independent variables (labeling method, normalization method, and feature extraction method) onto the cross-subject emotion recognition accuracy. The selected dataset for this analysis has been the SEED dataset, where the brainwaves of 15 participants were recorded while watching the same 15 film clips across three different sessions.

We first compared the Welch, multitaper, and differential entropy methods for extracting features in task of binary and ternary classification. Our participant-independent model was a CNN-based network with an input and three hidden layers each followed by our new, stratified normalization method, and an output followed by a softmax function for classification. The highest leave-one-out cross-validation mean accuracy with our model was M = 0.916, SD = 0.074 for binary classification and M = 0.796, SD = 0.104 for ternary classification, when extracting the features with the multitaper method. We also compared our stratified normalization method with batch normalization, obtaining after implementing an ANOVA test that the classification accuracies for stratified normalization was statistically higher than batch normalization. We also observed that including the neutral labels in the model hinders the classification of the negative labels, decreasing their classification accuracy from 90.67% to 75.11%.

Then, we found out that implementing stratified normalization is highly efficient in reducing the inter-participant variability from the data. Indeed, by training SVMs to try and recognize which participant the activation data of a given layer belongs to, we could observe that the participant identification information, or brain signature, was lost from a layer to another.

As we compared the embeddings at the level of the input and output layers, we could see that the stratified normalization already erases this brain signature in the input layer, such that by the end of the network, it is almost gone already – 33% for two categories and 31% for three categories in the last layer of the models with stratified normalization, approaching a chance level of 20%. It would be interesting to look for new ways of improving this result further.

Regarding the published articles, our method outperforms the rest of the proposed methods for binary classification and overpasses the works that did not use the data for validation during the training process for ternary classification.

These results indicate the high applicability of stratified normalization for cross-subject emotion recognition tasks, suggesting that this method could be applied not only to other EEG classification datasets but also to other applications that require domain adaptation algorithms.

## Footnotes

1 https://github.com/javiferfer/Cross-subject-EEG-emotion-recognition-through-NN

